# 3D-printed capillaric ELISA-on-a-chip with aliquoting

**DOI:** 10.1101/2022.09.23.508398

**Authors:** Azim Parandakh, Oriol Ymbern, Will Jogia, Johan Renault, Andy Ng, David Juncker

## Abstract

Sandwich immunoassays such as the enzyme-linked immunosorbent assay (ELISA) have been miniaturized and performed in a lab-on-a-chip format, but the execution of the multiple assay steps typically requires a computer or complex peripherals. Recently, an ELISA for detecting antibodies was encoded structurally in a chip thanks to the microfluidic chain reaction but the need for precise pipetting and intolerance to commonly used surfactant concentrations limited the potential for broader adoption. Here, we introduce the ELISA-on-a-chip with aliquoting functionality that obviates the need for precise pipetting, accommodates higher surfactant concentrations, includes barrier channels that delay the contact between solutions and prevent undesired mixing, and that executed a quantitative, high sensitivity assay for the SARS-CoV-2 nucleocapsid protein in 4×-diluted saliva. Upon loading the chip using disposable pipettes, capillary flow draws each reagent and the sample into a separate volumetric measuring reservoir for detection antibody (70 µL), enzyme conjugate (50 µL), substrate (80 µL), and sample (210 µL), and splits washing buffer into 4 different reservoirs of 40, 40, 60, and 20 µL. The excess volume is autonomously drained via a structurally encoded capillaric aliquoting circuit, creating aliquots with an accuracy of >93%. Next, the user click-connects the assay module, comprising a nitrocellulose membrane with immobilized capture antibodies and a capillary pump, to the chip which triggers the step-by-step, timed flow of all aliquoted solutions. A colored precipitate forming a line on a nitrocellulose strip serves as an assay readout, and upon digitization, yielded a binding curve with a limit of detection of 54 and 91 pg/mL for buffer and diluted saliva respectively, vastly outperforming rapid tests. The ELISA chip is 3D-printed, modular, adaptable to other targets and assays, and could be used to automate ELISA in the lab; or as a diagnostic test at the point of care with the convenience and form factor of rapid tests while preserving the protocol and performance of central laboratory ELISA.

## Introduction

The enzyme-linked immunosorbent assay (ELISA) is utilized for the detection and quantification of proteins, antibodies, or antigens. The sandwich format with a capture antibody immobilized on the surface and a detection antibody applied in solution is used for assays requiring high sensitivity and specificity. Laboratory microplate ELISA still serves as a gold standard for assays and benefits from high sensitivity thanks to enzymatic signal amplification (down to sub-picomolar concentration for the best antibody pairs), quantitative results, standardized operations, off-the-shelf consumables, and a comparably high throughput thanks to the use of 96-well plates. Long incubation times and copious washing between different steps to reduce non-specific binding and assay background are critical to achieving high assay sensitivity. However, the ELISA suffers from several downsides, such as being laborious, lengthy (∼2 - 12 h depending on the protocol), requiring precise timing for each step, dependence on technical skills notably for adding and removing reagents (and thus susceptible to inter-operator variation) and necessitating a plate reader for signal readout.^1,2^

The miniaturization of ELISA has proceeded thanks to microfluidic lab-on-a-chip systems that can also automate the protocol.^3,4^ Microfluidics successfully reduced the consumption of reagents and the total assay time while preserving assay performance. However, whereas the chips are small, they rely on bulky peripherals such as syringe pumps^3^ or control motors,^4^ and a computer or an instrument for operation.^5,6^ Capillary phenomena and gravity have been harnessed to automate simple liquid manipulation, reducing or obviating the need for an external/active power supply.^7–9^ For instance, a disk-like microfluidic platform (powered by a combination of centrifugal and capillary forces)^8^ and a microfluidic siphon platform (powered by gravitational forces)^9^ have been developed to carry out the common steps of a conventional ELISA with reduced reagents consumption and assay time while preserving assay sensitivity. Yet, both examples require multiple precise pipetting steps and timed user interventions for operation.

Sandwich assays can also be performed at point-of-care using so-called lateral flow assays (LFAs), also called rapid diagnostics, and are used globally for pregnancy tests and COVID-19 diagnosis. LFAs replace the enzyme amplification with conjugated colorimetric particles (either gold nanoparticles or polystyrene beads) that become visible to the naked eye upon accumulation. LFAs are simple to use as they only require the application of the sample, which flows thanks to capillarity without the need for peripherals, and produce a test result within a few minutes. However, LFAs offer only qualitative yes-no results, their sensitivity is typically lower compared to that of laboratory ELISA and are also not suitable for archival as the readout must be completed within a few minutes of the test, because otherwise, the result can be compromised.^10–12^ Enzymatic amplification has been implemented in the LFAs,^13,14^ for instance by using a microfluidic interface,^14^ to improve sensitivity. Yet, they cannot implement various fluidic handling tasks of common ELISA such as timed incubation of reagents and multiple rinsing steps between each incubation interval.

Paper-based microfluidics has been developed to introduce more advanced fluidic functions such as sequential delivery, additional rinsing step, and as well as enzymatic amplification that collectively help improve assay sensitivity compared to LFAs.^15–17^ Sponge actuators that upon swelling connect or disconnect different parts of a paper-based microfluidic circuit, along with flow paths with different lengths and resistance, have been used to time the delivery of multiple reagents for completing a bona fide ELISA.^17^ However, these systems lacked the intermediate washing steps characteristic of classical ELISA, and undesired mixing of consecutive reagents occur at their mutual interface. Enzyme-substrate mixing may limit the potential for higher sensitivity as it could contribute to non-specific signal amplification.

Capillaric circuits (CCs) are capillary microfluidics in microchannels designed and built using capillaric elements which can automate liquid handling operations by pre-programming them structurally using capillary phenomena and powering them by capillary flow, without the need for peripheral equipment.^18,19^ Multiple CCs have been designed to perform and automate ELISA with new functionality including flow reversal,^18^ timing, reagent lyophilization,^20^ and portable readers,^21,22^ but they skip intermediate washing steps. Aliquoting of a single solution into multiple reservoirs has been shown for a nucleic acid test.^23^ For an ELISA in a CC, multiple solutions including sample, buffer and reagents must be serially delivered, and fluidically connected to effect fluid flow by hydraulic transmission of pressure differentials, which would subject them to unwanted mixing due to diffusion and/or convection, negatively affecting assay sensitivity and reliability.

The microfluidic chain reaction (MCR) introduces conditional initiation of capillary flow events, whereby event *n* is triggered after the preceding event *n-1* has been completed, and completion of *n*, in turn, initiates event *n+1; the condition is encoded using the so-called capillary domino valves*.^24^ MCR can drive hundreds of sequential flow operations robustly, thus opening new opportunities for CCs. CCs and MCRs are susceptible to failure in the presence of surfactants that reduce surface tension and contact angles. Yet surfactants are essential ingredients to assays, and often 0.05% Tween 20 is used to prevent non-specific binding.^25^ In our previous MCR demonstrations, we could only accommodate 0.0125% because higher concentrations led to corner flow and trapping of air bubbles^19^ and failure of stop valves.

User-friendliness is one of the critical features for a device to be used at the point of need,^26–28^ and accurate aliquoting and volumetric consistency are essential to reliable immunoassays.^29,30^ Whereas in the lab they can be met using precision pipettes operated by technicians, they are difficult to achieve in a point-of-care setting. Both ELISA and the MCR CC introduced previously were dependent on precision pipetting. LFAs for COVID-19 (coronavirus disease 2019) often include a dropper and instructions for delivering a precise number of droplets.^31^ Droppers are prone to occasional miscounting,^31^ while for more complex assays multiple solutions with different volumes are required that could not be serviced using droppers to be supplied and aliquoted with different volumes.

Here, we introduce the ELISA chip that automates ELISA protocol on a chip using an MCR CC while preserving the washing steps used in classical ELISA. The ELISA chip circumvents the need for precise pipetting thanks to automated aliquoting of solutions and can accommodate higher surfactant concentrations commonly used in immunoassays. Akin to measuring spoons that are used to size ingredients in cooking, measuring reservoirs with different volumetric capacities are used to aliquot reagents, buffer, and sample. Upon loading, solutions spontaneously fill their respective measuring reservoir, while an integrated capillaric aliquoting circuit (CAC) autonomously drains excess liquid from all reservoirs simultaneously, forming accurate aliquots.

We describe the capillaric circuitry and its components, and the step-by-step automated capillary flow operations of both aliquoting of all solutions and the sequential MCR-controlled ELISA protocol. We characterize the volumetric accuracy of aliquoting, the timing precision of delivery of the reagents for the ELISA, and the performance and limit of detection (LOD) of an assay for the detection of the SARS-CoV-2 (severe acute respiratory syndrome coronavirus 2) nucleocapsid (N) protein spiked in buffer and 4×-diluted human saliva.

## Results and discussion

### ELISA chip

We designed a capillaric ELISA chip without a moving part that automates aliquoting of sample, reagents, and buffer, and autonomously executes an ELISA protocol by flowing eight solutions in sequence according to predetermined flow rates using an MCR, Fig. 1. The solutions are delivered for the ELISA in the order of (i) sample, (ii) buffer (iii) biotinylated detection antibody, (iv) buffer, (v) streptavidin poly-HRP (horseradish peroxidase), (vi) buffer, (vii) colorimetric substrate DAB (3,3′-Diaminobenzidine) and (viii) buffer, Fig. 1a. The ELISA was developed for measuring the SARS-CoV-2 N protein in natural saliva and includes a strip of nitrocellulose membrane with a positive control line and a test line that will produce a permanent colorimetric signal proportional to the concentration of the N protein.

**Fig. 1.**
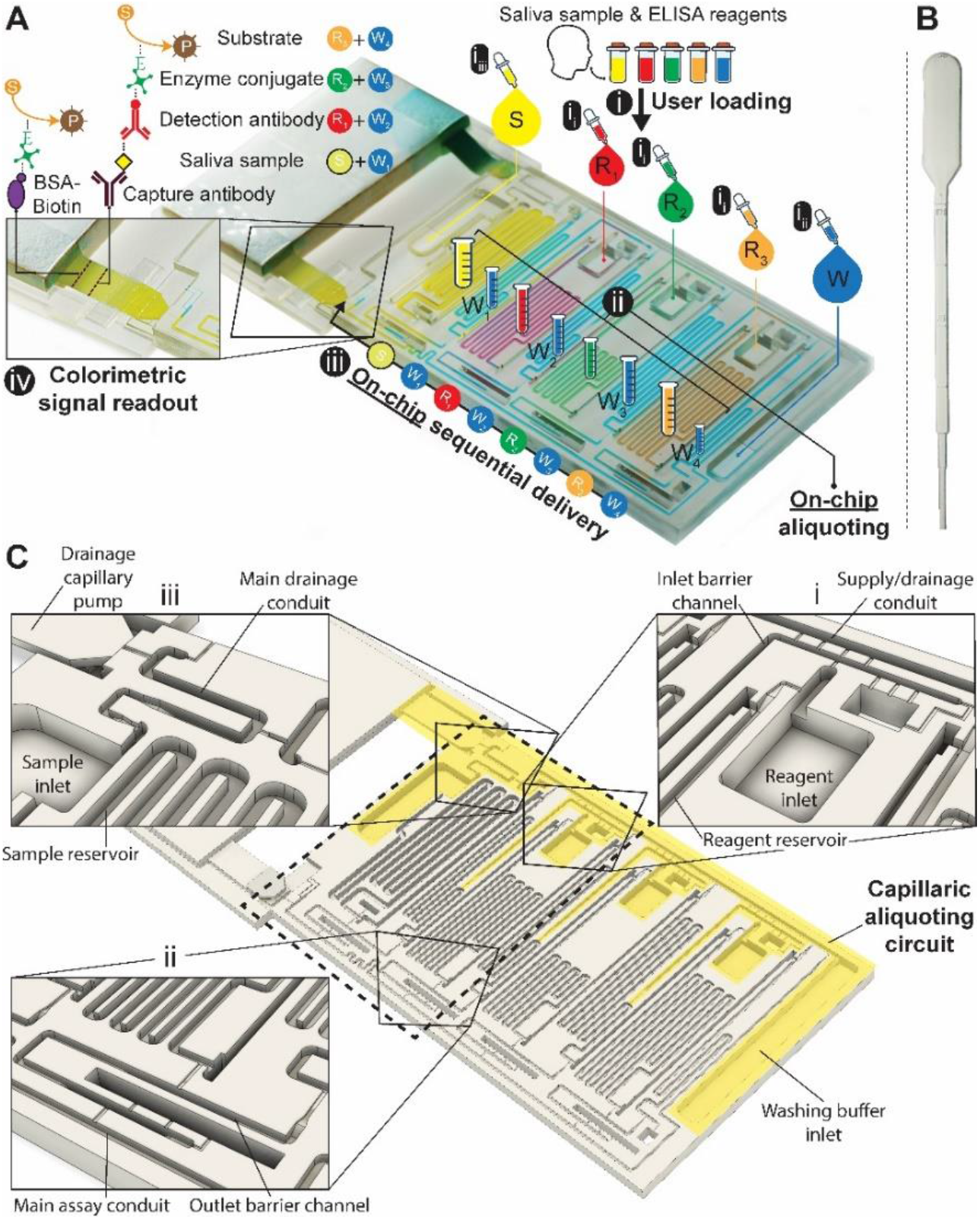
3D-printed ELISA chip with aliquoting for SARS-CoV-2 N protein assay in saliva. (A) Picture of the chip with superposed process flow graphics. (i_i_) Reagents (R_1-3_) and (i_ii_) washing buffer (W) are added to the inlets using disposable, low-precision pipettes shown in (B) and fill the measuring reservoirs by capillary flow; the washing buffer is split into four reservoirs automatically (W1-4). (i_iii_) Addition of the sample brings the drainage capillary pump to a fluidic connection, initiating drainage of excess solutions from all inlets via the capillaric aliquoting circuit (CAC, see C), and producing (ii) final aliquots schematized by graduated tubes. (iii) Connection of the nitrocellulose strip triggers the ELISA via the sequential flow of sample, reagents, and washing buffer through the chip according to the propagation of the MCR. (iv) Following the enzymatic conversion of the substrate into a permanent line of a brown precipitate, the result can be read out by eye, and quantified following digitization. (B) Picture of a disposable pipette used to load the chip. (C) 3D design view of the ELISA chip with the structurally encoded capillaric elements without moving parts. The CAC is highlighted in yellow, and insets showcase (i) inlet barrier channel and supply/drainage conduit and air vent, (ii) reservoir outlet, outlet barrier channel, main assay conduit, and air vent (iii) sample inlet and reservoir, main drainage conduit and connection to supply/drainage conduit, and drainage capillary pump.

The need for precise pipetting and the use of laboratory pipettes is circumvented by integrating CAC into the chip which accepts the delivery of larger-than-needed volumes into the adequately sized inlets, followed by the spontaneous flow of the solutions via capillarity into serpentine measuring reservoirs for each solution. To complete the aliquoting process, the CAC removes excess volumes into a drainage capillary pump. The volume of each reservoir was set following assay optimization.

Simple squeeze pipettes, Fig. 1B, can be used to load the chip while visually monitoring filling progression, and the measuring reservoirs form the aliquots (symbolized by graduated tubes) after draining excess liquid. The ELISA chip has five inlets servicing the eight reservoirs, three for the reagents, one for the buffer, and one for the sample, which need to be added in this order. The buffer is automatically split into four separate measuring reservoirs on the chip. The addition of the sample simultaneously fills the sample reservoir and initiates the drainage of excess volumes via the CAC. Next, upon click-connection of the main capillary pump, the step-by-step execution of the ELISA protocol is triggered, and the eight reservoirs chained by capillary domino valves are drained one by one as the MCR progresses. Finally, the signal is read out via the naked eye or using a scanner for quantification.

### Step-by-step loading and aliquoting

The ELISA chip loading and aliquoting are shown in Video S1. Using a low precision pipette (Fig 1B), the user first deposits each reagent in the respective inlet in an arbitrary order with an excess of solution. The three serpentine reservoirs for the detection antibody, streptavidin poly-HRP, and DAB solutions have nominal measuring volumes of 70, 50, and 80 µL, respectively. Fig 2 A & B show the loading of detection antibody and are representative of the other reagents. Capillary flow fills the reservoir up to the trigger valve at the empty outlet barrier conduit, while excess solution remains in the inlet and fills the trigger valve connecting to the conduit at the inlet (called inlet barrier channel) that is part of the CAC (See Fig. 1C). Next, the washing buffer is delivered to the common inlet to the right of the ELISA chip and flows by capillarity through the supply/drainage conduit into the four washing buffer measuring reservoirs with a nominal volume of 40, 40, 60, and 20 µL, Fig 2C. The buffer simultaneously fills the main assay conduit, which is supplied by the vertical channel adjacent to the buffer inlet. The buffer distribution and splitting simplify chip loading as only one buffer solution needs to be added. However, the chip design imposes contact between buffer and reagents both at the inlet and outlet of the reagent reservoirs which could lead to unwanted mixing by diffusion and convection between various solutions.

**Fig. 2:**
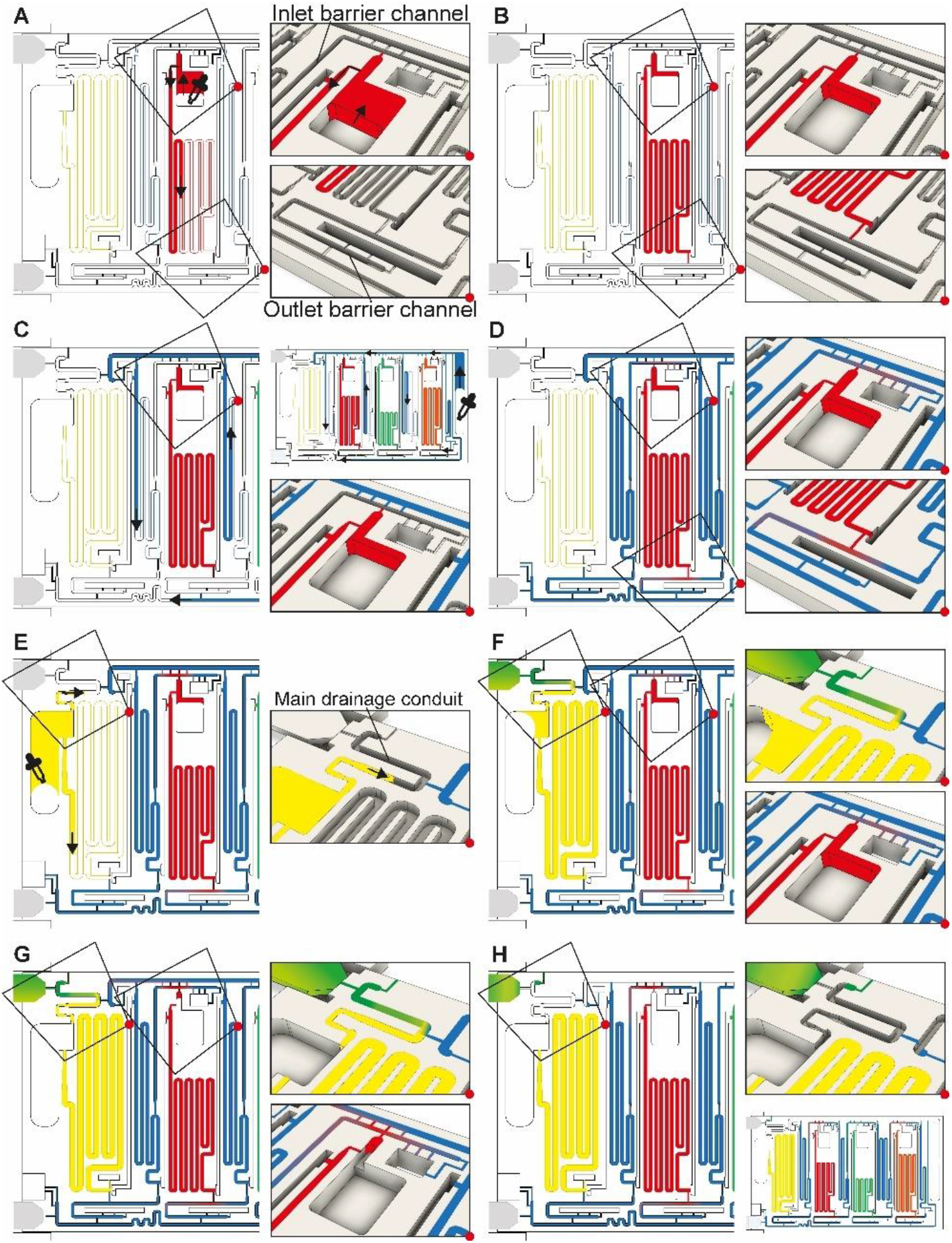
ELISA chip loading, filling into measuring reservoirs, and drainage of excess liquid by the CAC. Most figures show the part of the chip outlined by the dashed rectangle in Fig 1C. (A & B) Loading of the detection antibody reagent into the inlet followed by capillary flow into the measuring reservoir and stop of the liquid at the capillary stop valves located next to the inlet and at the outlet. Loading of the other two reagents, i.e., streptavidin poly-HRP and DAB, follows the same procedure. (C & D) Loading of buffer in buffer inlet and splitting into the buffer reservoirs supplied by the supply/drainage conduit. (E-H) Sample loading with the filling of the sample reservoir and simultaneous triggering of the drainage of all excess solutions on the chip via the CAC. See Video S1 and text for further details.

### Suppressing unwanted mixing and bubble trapping

We isolated each of the reagents measuring reservoirs to prevent unwanted mixing between them and/or the sample prior to assay completion. We equipped them with two diffusion barrier channels located upstream (at the inlet) and downstream (at the outlet) of each reagent reservoirs. By isolation, we suppress undesired preliminary mixing of reagents that could affect assay performance and reproducibility during both aliquoting (i.e., filling and excess volume drainage), and subsequent ELISA sequential delivery (e.g. premature reaction between enzyme and colorimetric substrate DAB could lead to unwanted background signal). Because diffusion will occur for a long time at the outlet of each reservoir, and for almost the entire assay duration particularly for the DAB solution, the last reagent to flow in the ELISA protocol, a long diffusion barrier is needed. By implementing diffusion barrier channels, we increase the distance between ELISA reagents to prevent premixing that could occur by both diffusion and/or convection (e.g. small convective flows due to evaporation). The outlet barrier channel was designed with a U-turn and a separation length between the outlet of the reservoir and the main assay conduit with a length of 13-17 mm. The characteristic one-dimensional diffusion time is given as t= L^2^/D with L = length, D = diffusion constant and t = time.^32^ For an IgG antibody with D = 35 µm^2^/s and DAB with D = 550 µm^2^/s, the time for diffusion across 10 mm is ∼16.5 days and ∼1 day, respectively. In our case the functional connection connecting the reservoir and the barrier forms a constriction,^24^ hence the diffusion time would be further increased relative to the calculation.

The inlet and outlet barrier channels form a separation between each reagent reservoir and the supply/drainage conduit (upstream), and main assay conduit (downstream) respectively. Both barrier channels remain empty during autonomous filling of the reagents due to the action of trigger valves (see Fig 2A), and autonomously fill by buffer after buffer injection, (Fig 2D). The inlet of the inlet (upstream) barrier conduit is located to the left of the reagent reservoir and branches out from the serpentine buffer reservoir, close to the outlet of the buffer reservoir. Upon reaching the branching point, capillary flow filling the buffer reservoir immediately branches into the barrier channel and fills it up to the extremity, which forms a dead-end. To prevent bubble trapping, an air vent is included that is connected to the channel via four stop valves, including three positioned upstream of the dead-end. At the reservoir outlet, the outlet barrier channel, which forms an extension to the buffer reservoir, is similarly filled by the buffer. However, the buffer reservoir to the right of the reagent channel supplies the corresponding outlet barrier channel and fills only after the buffer reservoir is filled completely (Fig 2D bottom close-up, Video S1). Akin to the inlet barrier channel, the extremity of the outlet barrier channels also forms a dead-end and is connected by three stop valves to a venting opening. Note that the design ensures that only pristine buffer fills each of the buffer measuring reservoirs. The inlet and outlet barrier channels are empty as the buffer starts flowing, and hence there is no contact or mixing between the reagents and the buffer flowing into the reservoirs.

The provision of multiple venting connections/stop valves is needed to accommodate liquids with low surface tension such as buffer containing a surfactant that is used in ELISA (e.g., 0.05% Tween 20) and other assays. Low surface tension liquids have a lower contact angle than water which has a high surface tension, which induces corner flow preceding the main filling front. The corner flow reaches the extremity of the conduit before the main filling front and can thus clog small air vents located at the extremity of dead-end channels before all the air can escape (See Supplementary Fig S1). Upstream connections to the vent allow air to escape even after corner flow reaches the dead-end and here prevent air bubbles from being trapped.

### Excess volume draining

Drainage of the excess volumes is initiated by supplying the sample to the sample inlet, Fig. 2F. The sample inlet is connected to both the sample measuring reservoir and the main drainage conduit, both filling simultaneously by capillary flow. The outlet of the sample reservoir connects to a now pre-filled outlet barrier channel supplemented with an air vent to permit complete filing. The sample flowing into the main drainage conduit also initiates drainage of the excess volumes by triggering a trigger valve connecting the filled supply/drainage conduit to the main drainage conduit, and subsequently flowing to the drainage capillary pump, all part of the CAC. Initially, most of the drainage flow is supplied by the sample owing to excess solution in the sample inlet and the absence of a retention capillary pressure, and lower flow resistance. Once the sample inlet is emptied to the point where only corners remain filled, the drainage flow rates from the inlets containing excess buffer and reagents increase. The inlet barrier channel, in addition to minimizing the diffusion between the main drainage conduit and each reservoir, also forms the fluidic connection through which the excess reagents is drained from each reagent inlet.

### Flow path routing during drainage

Whereas conceptually simple, in CCs, draining must both be triggered and stopped without external intervention. In addition, multiple possible drainage paths exist when the circuit is operating in the absence of active valves to close off sections of the circuit, yet to avoid unwanted mixing and drainage of solutions in the reservoirs, the liquid flow must flow through the pre-designed flow path, Fig. 3A. In the desired path, the excess reagents flow from the common inlet to the drainage capillary pump as follows: reagent inlet → inlet barrier channel → supply/drainage conduit → main drainage conduit → drainage capillary pump. To facilitate flow and favor this fluidic path while preserving other capillaric functionality, we included 4 parallel connection conduits (0.1×0.2×0.75 mm^3^) between the inlet barrier channel and the supply/drainage conduit (Fig. 1C close-up i), which yielded an overall flow resistance of 135 Pa·s·mm^-3^ for path i.

**Fig. 3.**
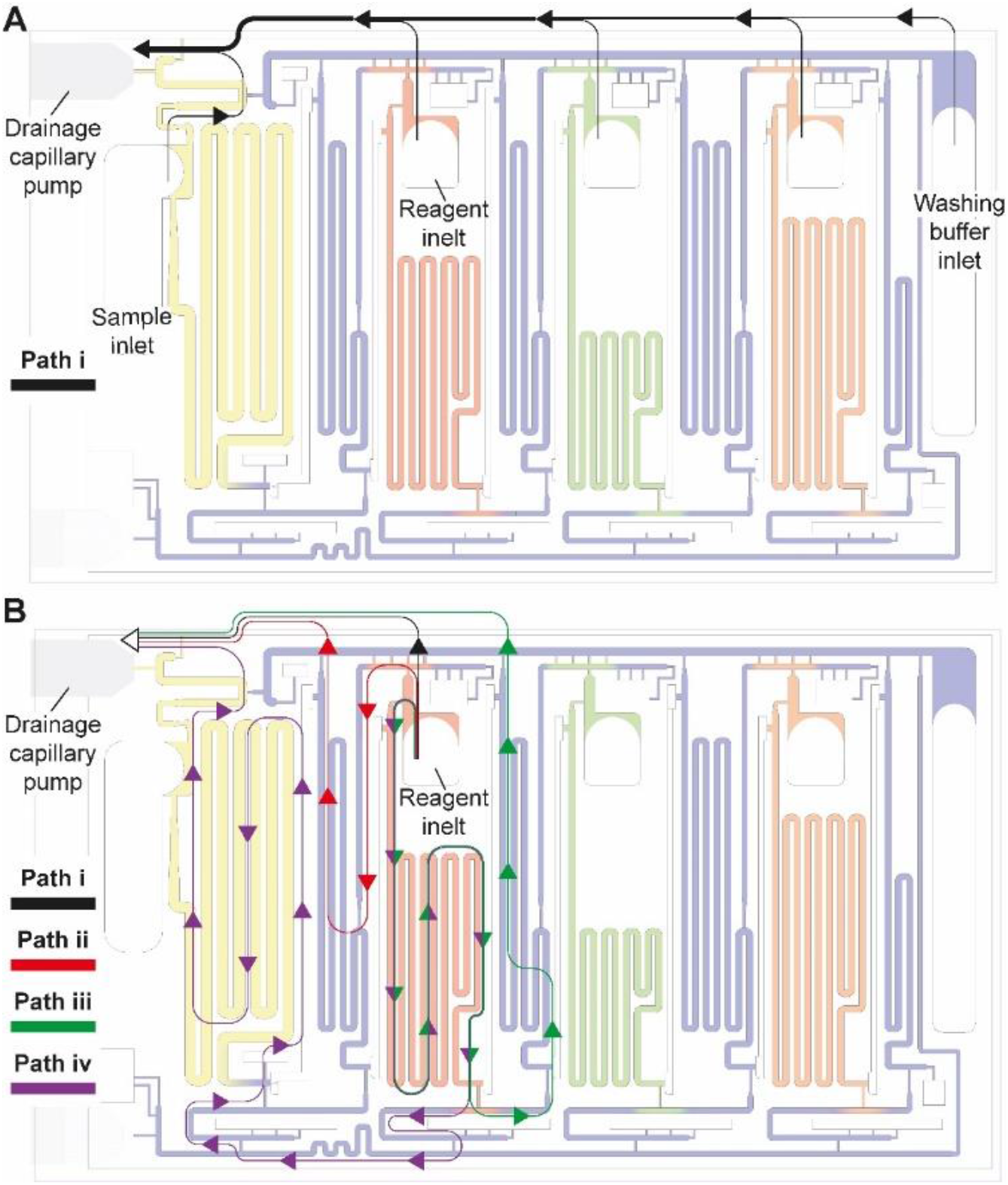
Drainage of excess volumes via the CAC. (A) The flow path for draining excess volumes in sample, reagent, and buffer inlets is shown with the black arrows. (B) In addition to the desired flow path i, three alternative, parasitic paths ii, ii and iv are schematized using colored arrows. Drainage of reagent via a parasitic flow path could lead to mixing with other solutions and jeopardize the function of the ELISA chip. The design minimizes parasitic flow via a design that ensures that their flow resistance is much higher because they are longer and if necessary, sections with a small cross section (and therefore a high flow resistance) are included. Note that in panel B, all flow paths are shown for only one reagent, and the same exist for other reagents as well.

For each of the three reagent inlets, three unwanted, leaky flow paths exist, as illustrated for the detection antibody in Fig. 3B. Flow through any of the parasitic paths should be disabled as they would result in unwanted reagents mixing and/or deviation from the pre-programmed volumes. Path ii proceeds from reagent inlet → inlet barrier channel → washing buffer reservoir → supply/drainage conduit, and path iii from reagent inlet → reagent reservoir → reagent outlet barrier channel → washing buffer reservoir → supply/drainage conduit to the main drainage conduit and drainage capillary pump. Both paths would result in reagents flowing into, and mixing with, the washing buffer in the adjacent reservoirs. To impede drainage through path ii, the resistance at the inlet of the inlet barrier channel as well as that of the corresponding washing buffer reservoir was increased, resulting in overall flow resistance of 250 Pa·s·mm^-3^ for path ii. Likewise, to impede drainage through path iii, the resistance at the inlet of the outlet barrier channel as well as that of the corresponding washing buffer reservoir was increased, thereby the overall resistance of path iii was 550 Pa·s·mm^-3^.

Path iv follows the reagent inlet → reagent reservoir → reagent outlet barrier channel → main assay conduit → sample outlet barrier channel → sample reservoir → sample inlet → main drainage conduit → capillary drainage pump (Fig. 3B, path iv), with an overall resistance of 700 Pa·s·mm^-3^. This drainage path would lead to mixing with the sample, yet the last part of being in charge of the excess drainage of the sample. This path only exists until the point where the excess sample is drained into the main drainage conduit (with an overall resistance of 50 Pa·s·mm^-3^) and is disconnected from the sample reservoir as the inlet is emptied.

Here we discussed the overall flow resistance of each of the four paths, but the parasitic flow is governed by the difference between the point where the parasitic paths split from the drainage path i to the point where they merge again, which is a shorter distance. The ratio of flow resistance for the split section between path i, and each of paths ii, iii, and iv was calculated to be 7.8, 8.4, and 6.9 times higher respectively. Path iv is only active for a short time, and thus only contributes marginally. This design hence ensures that most of the excess volume will flow through path i.

### The automated ELISA protocol

The ELISA is initiated by the user click-connecting the main capillary pump to the chip (Fig. 4A), which wicks liquid from the CC and the reservoirs, and initiates the MCR that propagates from one reservoir to the next via the capillary domino valves that chain the reservoirs,^24^ (movie S2). Briefly, the first reservoir is the sample reservoir that is capped by a retention burst valve (RBV) with a nominal capillary/burst pressure of -200 Pa (i.e., it will burst when the negative sucking pressure in absolute value is >200 Pa), which quickly bursts and results in draining of the reservoir. Air drawn in from the outside replaces liquid, and once all liquid is displaced, the capillary domino valve that forms an air link to the next reservoir is now connected to the outside via the just-emptied reservoir. At the top of the buffer reservoir, there is another RBV (here with the nominal capillary pressure of -215 Pa), which in turn bursts, leading to the draining of the first buffer reservoir, and in turn, opening the air link to the next (reagent) reservoir, and so on. The sequence of flow is thus sample, buffer, detection antibody, buffer, streptavidin poly-HRP, buffer, DAB, and buffer (Fig. 4B-F). The initiation of draining of the main assay conduit is not controlled via the MCR because the chip lacked space for an additional capillary domino valve. Instead, we implemented an additional RBV with a nominal burst pressure of -550 Pa. That is the RBV with the highest threshold within the CC, and hence the last one to burst. The ELISA chip illustrates the possibility of combining an MCR with an RBV to control the sequence of liquids flowing through an assay conduit.

**Fig. 4.**
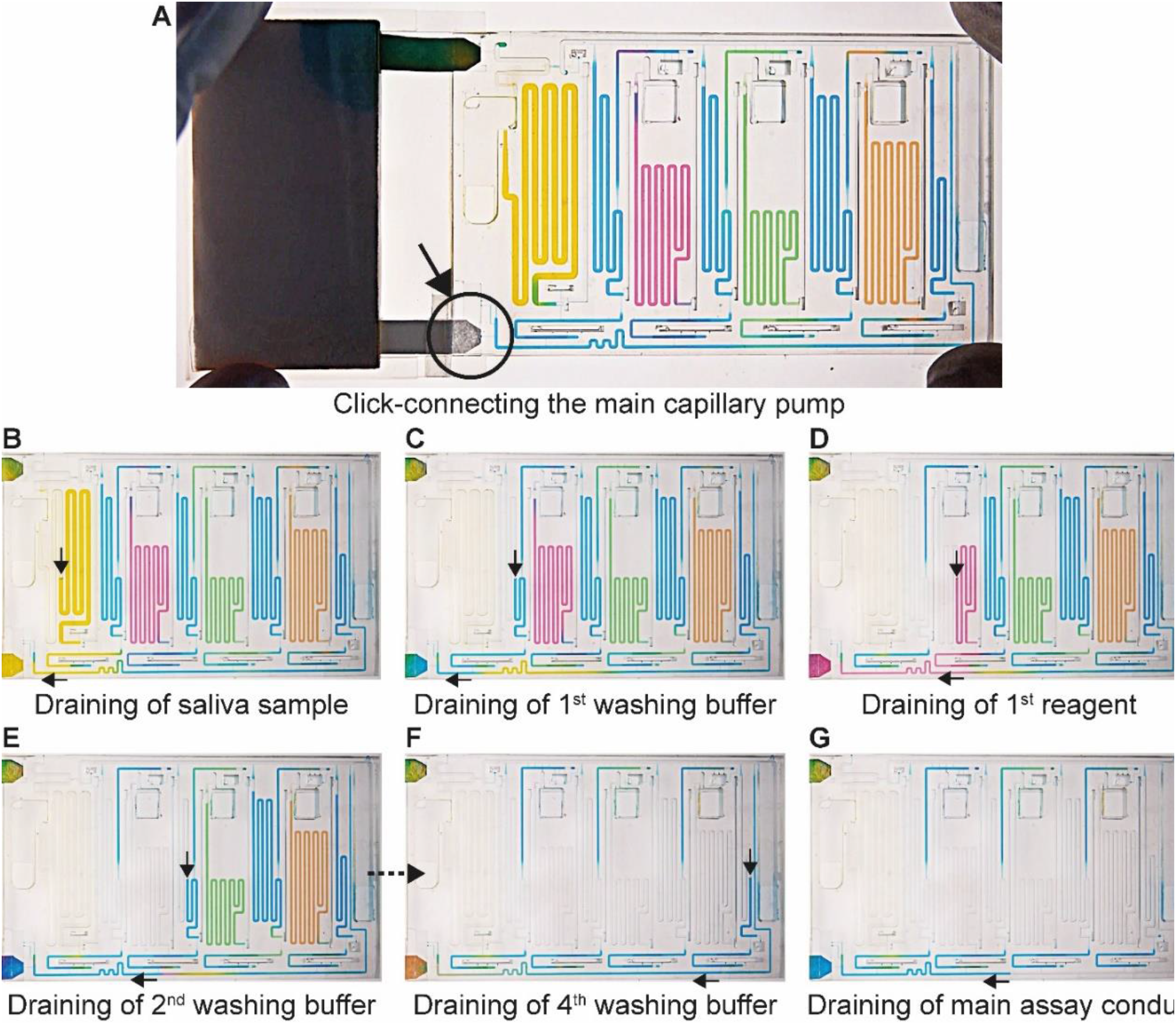
MCR progression of the ELISA chip to complete ELISA steps. (A) Click-connecting the main capillary pump to the chip (the region indicated by the black circle and arrow) triggers (B-G) sequential delivery of the sample, reagents, and corresponding washes to the sensor. Black arrows indicate the direction of flow.

**Fig. 5.**
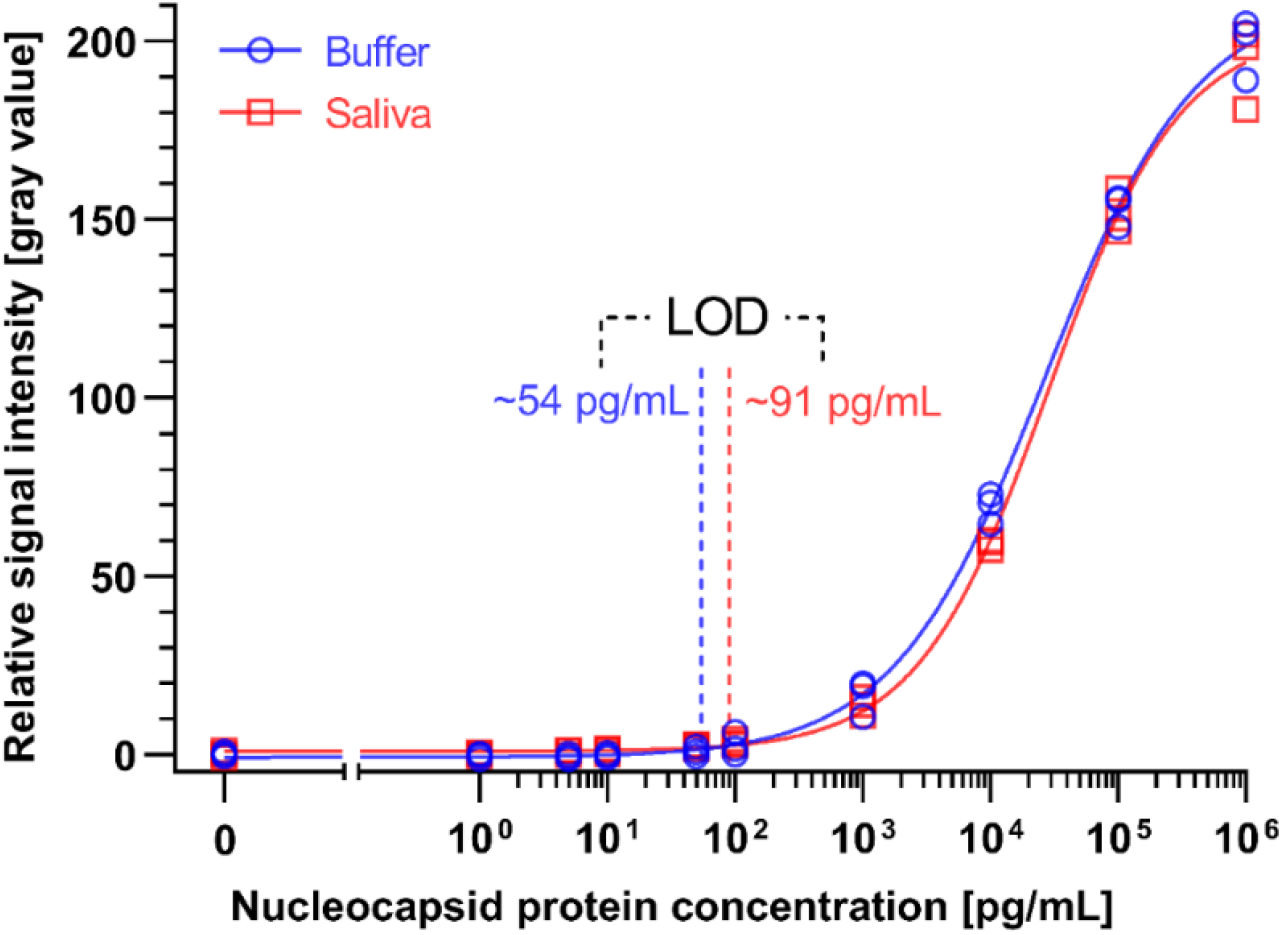
Calibration curves of the SARS-CoV-2 N protein assay enabled by the ELISA chip. Different concentrations of SARS-CoV-2 N protein were spiked in ELISA buffer or 4×-diluted pooled saliva. The resultant time-insensitive colorimetric signals were then captured by a regular scanner and yielded a binding curve that was fitted with a 4-parameter logistic regression. The number of replicates for each concentration is 3 for each calibration curves.

### ELISA chip over-loading capacity and overflow

We evaluated the volumetric operational window of the ELISA chip that preserves aliquoting and overall ELISA chip functionality. We analyzed the video of four ELISA chips loaded using precision pipettes with known volumes of sample, reagents, and buffer, determined the minimum volume required to fill the measuring reservoirs, tested a range of excess volumes, and verified flow functionality while also monitoring unwanted mixing.

The nominal aliquot volume corresponds to the capacity of the measuring reservoir. However, in practice, some additional volume is needed to account for dead volumes in the chip. Conversely, if the inlets are overfilled, leakage into other parts of the chip could occur. We verified the nominal aliquoting accuracy by mapping the levels of the measuring reservoirs filled with food dyes after aliquoting was completed, and calculating the volumetric error. For the sample, the nominal aliquot volume is 210 µL but at least 300 µL is required because the sample triggers the CAC (as explained above). As this ELISA chip is designed for saliva, which was collected in an amount of 1 mL per individual, the additional volume requirement could be accommodated easily. We verified that the chips preserved their functionality and nominal aliquoting accuracy for a volume of 400 µL (maximum tested). Within the 300-400 µL range, the nominal accuracy of sample aliquoting was 99.5% within the coefficient of variation (CV) of 1.1%.

In the case of detection antibody, streptavidin poly-HRP, and DAB, the nominal aliquot volume is 70, 50, and 80 µL respectively, while at least an extra ∼1 µL (for a total of 71, 51, and 81 µL, respectively) is needed to completely fill each reservoir. The maximum volumes that reagent inlets could accommodate while avoiding pre-mixing of reagents with buffer were 110, 90, and 120 µL for the detection antibody, streptavidin poly-HRP, and DAB, respectively. Under these conditions, the nominal accuracy of aliquoting was found to be 99.7 (CV= 2.1%), 93.4 (CV= 1.5%), and 99.9% (CV= 3.5%) respectively. In the case the overloading exceeds the maximum volumes, the chip will continue to function, but some part of the excess may spill into other reservoirs, which was observed when loading 140 µl of detection antibody that led to spilling into the adjacent buffer reservoir, Fig. S3.

For the washing buffer, ∼300 µL is needed to fill all washing buffer reservoirs as well as the supply/drainage conduit, inlet/outlet barrier channels, and main assay conduit. We tested its operation for a volume up to 400 µl, successfully. Within this operating range, the buffer was reliably aliquoted into the four reservoirs of 40 µL, 40 µL, 60 µL, and 20 µL with a nominal accuracy of 98.7, 97.4, 98.3, and 94.7%, respectively, all within the CV of <1%. The results are summarized in Table 1. Considering these values, the ELISA chip outperforms the bulky pipetting robots utilized in laboratory ELISA, and indicates superior performance over the previously developed microfluidic devices while requiring significantly less user intervention.^29,30^

**Table 1.** ELISA chip over-loading, aliquoting, and flow timing performance.

|  | Aliquoting<sup>1</sup> |  |  |  | Sequential delivery<sup>1</sup> |
| --- | --- | --- | --- | --- | --- |
|  | Nominal aliquot volume (μL) | Loading range<sup>2</sup> (μL) | Visual aliquoted volume (μL)<sup>3,4</sup> | Nominal aliquoting accuracy (%) | Duration (min'sec'')<sup>4</sup> |
| **Sample** | 210 | 300-400<sup>6</sup> | 209.0 ± 2.4 (1.1%) | 99.5 | 24'39'' ± 40'' (2.7%) |
| **Detection antibody** | 70 | 71-110 | 70.2 ± 1.5 (2.1%) | 99.7 | 8'52'' ± 18'' (3.4%) |
| **Enzyme conjugate** | 50 | 51-90 | 53.3 ± 0.8 (1.5%) | 93.4 | 6'29'' ± 10'' (2.5%) |
| **Substrate** | 80 | 81-120 | 80.1 ± 2.8 (3.5%) | 99.9 | 10'57'' ± 22'' (3.4%) |
| **Wash #1** | 40 | 300-400<sup>6</sup> | 39.5 ± 0.1 (0.3%) | 98.7 | 4'58'' ± 7'' (2.4%) |
| **Wash #2** | 40 |  | 39.0 ± 0.3 (0.8%) | 97.4 | 5'2'' ± 6'' (2.0%) |
| **Wash #3** | 60 |  | 59.0 ± 0.2 (0.3%) | 98.3 | 7'57'' ± 12'' (2.5%) |
| **Wash #4** | 20 |  | 19.0 ± 0.3 (0.2%) | 94.7 | 3'11'' ± 6'' (3.1%) |

<sup>1</sup>Number of chips tested: 4
<sup>2</sup>Loading range that can be accommodated without impairing measuring function.
<sup>3</sup>Volumes are measured by visual analysis of liquid within the chip and calculating volume based on 3D design (see Experimental for detailed explanation).
<sup>4</sup>Reported in Mean ± Standard Deviation and CV in parenthesis
<sup>5</sup>Nominal aliquoting accuracy (%) = $(1 - \frac{|\text{Visual aliquoted volume} - \text{Nominal aliquot volume}|}{\text{Nominal aliquot volume}}) \times 100$
<sup>6</sup>Maximum volume tested

**Table 2.** **Figure of merit of the ELISA chip in comparison to a commercial microplate ELISA kit (SARS-CoV-2 N Protein Detection ELISA Kit from SinoBiological, Inc.) and an LFA rapid antigen test (R-Biopharm RIDA QUICK SARS-CoV-2 Antigen).**

|  | **Sino Biological SARS-CoV-2 N Protein Detection ELISA Kit** | **R-Biopharm RIDA QUICK SARS-CoV-2 Antigen Test** | **SARS-CoV-2 N Protein assay-on-ELISA Chip** |
| --- | --- | --- | --- |
| **Enzymatic amplification** | Yes | No | Yes |
| **Results type** | Quantitative | Qualitative (yes/no) | Qualitative and quantitative |
| **Readout timeframe** | Within 20 min after assay (= temporary) | Within 15 min after assay (= temporary) | No limitation (= permanent) |
| **Assay duration** | ~5 h | ~20 min | ~1.5 h |
| **LOD** | 35 pg/mL | 2,500 pg/mL<sup>1</sup> | 91 pg/mL |
| **Sample type** | Serum | PBS<sup>1</sup> | Human saliva |
| **Results interpretation** | Plate reader | Direct reading | Direct reading or scanner |
| **Precise pipetting** | Required | Required | Not required |
| **Hands-on time** | ~ 1 h | ~ 1-5 min | ~ 2 min |

<sup>1</sup>PBS = Phosphate buffer saline
<sup>2</sup>This value is the minimum concentration that could be detected by the naked eye, which is not equivalent to the LOD, a formulaic value.

### Flow timing

We characterized the reliability of the timing of the ELISA protocol by measuring the duration of each step pre-programmed into the ELISA chip. The timing of the assay steps is an important parameter when considering the accuracy and reproducibility of ELISA. We analyzed the video of four ELISA chips loaded with assay reagents spiked with food dye colors for visualization purposes. Table 1 summarizes the draining/delivery duration for the sample, reagents, and buffer. Remarkably, we found the timing to be reproducible within the CV of 3.5%. We consider these results, both for the volumes and incubation times, excellent for a capillary-driven microfluidic system and suitable for conducting ELISA.

### SARS-CoV-2 N protein ELISA

To evaluate the performance of the ELISA chip, we used it to run a saliva-based SARS-CoV-2 N protein assay. We performed an in-depth optimization process including testing various concentrations of capturing antibody, detection antibody, streptavidin poly-HRP, and Tween 20 as well as the extent of saliva dilution on the minimum detectable signal and assay background (See Table S1 for a summary of the optimization process).

Akin to classical microplate ELISA, the ELISA chip allowed for the implementation of enzymatic amplification that necessitates a two-step process (i.e., the addition of enzyme, followed by the substrate) which is not possible for the common LFAs that are typically carried out in one-step. DAB is the substrate used here which is chromogenic and oxidized in the presence of HRP, forming a brown precipitate at HRP locations. Whereas in laboratory ELISA the assay produces a colored solution, here it forms a precipitate on the paper strip which can be read out by the naked eye or digitized with a scanner^24^. Although the read-out time for both ELISA and LFAs is limited to a window of a few minutes, the precipitate in the ELISA chip is stable and can be read out later, thus potentially also serving as an archival record.

We generated binding curves by spiking-in N protein of SARS-CoV-2 across 6 orders of magnitude of dilution from 1 to 10^6^ pg/mL in 4×-diluted pooled human saliva and ELISA buffer. Fitting the experimental data using a 4-parameter logistic regression, we obtained an LOD of 54 pg/mL and 91 pg/mL for the N protein in buffer and 4×-diluted pooled saliva, respectively (Fig. 4). The difference in LOD between buffer and saliva is due to the use of saliva as the diluent in the saliva binding curve, which leads to a higher background signal and greater variation than buffer alone. The small dilution of human saliva together with the high sensitivity of the assay is a practical advantage of the developed ELISA-on-chip given the higher performance needed for reliable COVID-19 diagnosis based on saliva testing ^33^.

We benchmarked the ELISA chip against two commercially available microplate-based ELISA for SARS-CoV-2 N protein detection (SinoBiological, Inc. and RayBiotech Life, Inc.). These ELISA kits use blood serum as the sample and have a time-to-result of ∼5 h with an LOD of 35 pg/mL and 70 pg/mL, respectively, as reported by the manufacturer’s protocol. Note that the LODs of these serum ELISA were established based on a binding curve using buffer, and are thus comparable to the ELISA chip LOD for buffer; besides, RayBiotech calculated the LOD based on 2×SD above the blank instead of the widely used 3×SD. The LOD of the ELISA chip rivals one of the classical ELISA kits while being four times faster with less hands-on time and no time-sensitive manipulations of liquids, and in a format that is compatible with the point-of-care setting.

A study compared the performance of seven LFA rapid antigen tests for N protein spiked in phosphate buffer saline.^34^ The tests notably included the widely used Abbott Panbio COVID-19 Ag Rapid Test and the Roche-SD Biosensor SARS-CoV Rapid Antigen Test. The most sensitive one was reported to be the R-Biopharm RIDA QUICK SARS-CoV-2 Antigen Test which yielded a line visible to the naked eye for a concentration as low as 2.5 ng/mL.^34^ The LOD of the ELISA chip is ∼50 and ∼25 times higher than this LFA test in buffer and 4×-diluted pooled saliva, respectively.

### Conclusions and future works

We presented an integrated ELISA chip that miniaturized and automated an ELISA-on-a-chip using capillarics and an MCR with an encoded aliquoting function, enabling the ELISA chip to be serviced with disposable pipettes. The ELISA chip generated aliquots with various volumes and timed the assay steps both within the CV of ≤ 3.5%, rivaling the common pipetting robots utilized in laboratory ELISA and other automated microfluidic ELISA.^29,30^ The ELISA chip could operate with 0.05% Tween 20 commonly used in assays. The LOD of the ELISA chip for the SARS-CoV-2 N protein in 4×-diluted saliva was 91 pg/mL, in line with classical microplate ELISA and outperforming conventional lateral flow assays by ∼25×.

The ELISA chip could be 3D-printed and assembled in less than 1 h, and ∼1200 chips were manufactured as part of this work. ELISA chips were designed with superficial channels only and may thus be adaptable to mass production by injection molding with much lower mass manufacturing costs than 3D printing.

In the future, the chip may be validated with patient samples in retrospective and possibly prospective studies. It would be desirable to further simplify the operations and speed up the time-to-result to match LFAs, which could be facilitated by pre-drying reagents in the chip and rehydrating them with buffer.^20–22^ Following these improvements, ELISA Chips could be deployed at the point-of-need and used by non-experts, and using a cell phone for imaging and quantifying the assay results,^21,22,24^ quantitative, point-of-care test with the performance of a central laboratory ELISA become available for everyone.

## Experimental

### ELISA chip fabrication and preparation

The chips were designed in AutoCAD (Autodesk), exported as “STL” files, and printed with a stereolithography 3D printer with the LED wavelength of 405 nm (Pr 110, Creative CADworks, Concord, Canada) using a monocure 3D rapid clear resin (Monocure 3D, NSW, Australia) with the following printing parameters: exposure time per layer: 2.5 s (10 s for the base layer); transition buffer layers: 2; layer thickness: 20 µm; printing delay: 1 min; and gap adjustment: 0.1 mm. Once printed, the chips were cleaned with isopropanol (Fisher Scientific, Saint-Laurent, Quebec, Canada), dried under a stream of pressurized nitrogen gas, cured for 1 min in a UV lamp (CureZone; Creative CADWorks; Concord; Canada), plasma treated for 10 sec at 100% power (PE50 plasma chamber, Plasma Etch, Carson City, USA), and sealed with a microfluidic diagnostic tape (catalog number: 9795R; 3M Science. Applied to Life.™, Ontario, Canada).

A strip of Whatman CF4 paper (Cytiva, Marlborough, Massachusetts, United States) was clamped between 2 absorbent pads (Electrophoresis and Blotting Paper, Grade 320, Ahlstrom-Munksjo Chromatography, Helsinki, Finland) from the back end to collectively serve as the capillary pump. For the main capillary pump, a strip of Vivid™ 120 lateral flow nitrocellulose membrane (Catalog number: VIV1202503R; Pall Corporation, Port Washington, USA) was clamped between the same absorbent pads from the back end and to a G041 glass fiber conjugate pad (Millipore Sigma, Oakville, Ontario, Canada) from the front end to facilitate connection to the chip.

### Nitrocellulose membrane

The strips of vivid™ 120 lateral flow nitrocellulose membranes were designed in AutoCAD with the dimensions of 5.2 mm wide and 12 mm long and cut using the Silhouette Portrait paper cutter (Silhouette, Lindon, USA). Membranes were stripped with a 5 mm-wide test line of SARS-CoV-2 N protein mouse monoclonal antibody (catalog number: 40143-MM08; Sino Biological, Inc., Beijing, China) at the concentration of 1 mg/mL and a 5 mm-wide control line of bovine serum albumin (BSA)-biotin solution at the concentration of 50 µg/mL, both delivered using a programmable inkjet spotter (sciFLEXARRAYER SX, Scienion, Berlin, Germany). The membranes were dried for 1 h at 37 °C and blocked by dipping into the blocking buffer solution (1% BSA and 0.1% Tween 20 in PBS) until completely wet, followed by shaking on a rocker for 60 min at 75 rpm. The membranes were then retrieved, dried in an oven for 1 h at 37 °C, and stored with a desiccant at 4 °C until use on the next day.

### SARS-CoV-2 N protein ELISA

The sample solutions were prepared by spiking SARS-CoV-2 N protein (catalog number: 40588-V08B; Sino Biological, Inc., Beijing, China) at the concentrations of 0, 1, 5, 10, 50, 10^2^, 10^3^ 10^4^ 10^5^, 10^6^ pg/mL in either the ELISA buffer solution (0.1% BSA and 0.05% Tween 20 in PBS) or 4×-diluted pooled saliva solution. Fresh saliva specimens were collected using oral cotton swabs (Salivette, Sarstedt, Numbrecht, Germany), pooled, filtered through a 0.22-micron filter, and diluted by a factor of 4 in the ELISA buffer solution. The biotinylated SARS-CoV-2 N protein rabbit monoclonal antibody (catalog number: 40143-R004-B; Sino Biological, Inc.; Beijing, China) and streptavidin poly-HRP (Pierce; catalog number: 21140; ThermoFisher; Ottawa, Canada) solutions were prepared in the ELISA buffer solution both with the concentration of 7.5 µg/mL. The substrate solution was prepared by dissolving SIGMAFAST™ DAB tablets (catalog number: D4293-50SET; Sigma-Aldrich; Oakville, Canada) in Mili-Q water. The washing buffer solution was the same as the ELISA buffer solution.

For benchmarking, the developed SARS-CoV-2 N protein assay was compared with the SARS-CoV-2 (2019-nCoV) Nucleocapsid Detection ELISA Kit (catalog number: KIT40588; Sino Biological, Inc.; Beijing, China) and the RayBio® COVID-19 / SARS-COV-2 Nucleocapsid Protein ELISA Kit (catalog number: ELV-COVID19N; RayBiotech Life, Inc.; Peachtree Corners, United States).

### Nitrocellulose membranes image analysis and LOD calculation

After completion of the ELISA, the nitrocellulose strips were removed from the ELISA chip, left to dry at room temperature, and scanned at 1200 dpi in TIFF format (Epson Perfection V600) (see Supplementary Fig. S4). The images were imported in Photoshop (Version: CS5 ME) and superposed with guide structures to locate the region of interest (2.5 × 0.4 mm) for the test line as well as the bottom background and top background, each located 1.5 mm below and above the test line respectively. The superposed images were then imported in Fiji to measure the gray value of the three regions of interest for each nitrocellulose membrane (see Supplementary Fig. S5). For each concentration and the negative control, the local signal intensity of the test line was calculated by subtracting the gray value of the test line from the average gray value of the top and bottom local backgrounds. The relative signal intensity was then calculated by subtracting the local signal intensity of the test line from the average of the local signal intensity of the negative controls.

The experimental data were fitted using a 4-parameter logistic regression with the following equation:^35^

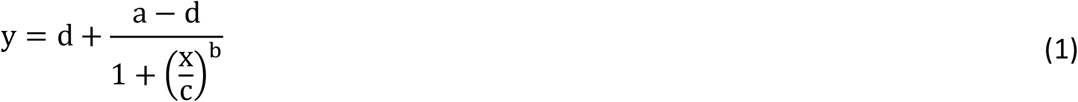

Where a and d are theoretical responses at zero and infinity, respectively, b denotes the slope factor (i.e., Hill slope), and c represents the mid-point concentration (inflection point).^35^

The LOD was then determined by adding 3 standard deviations to the mean relative signal intensity of the bank samples (i.e., zero antigen concentration) and calculating the corresponding concentration from the established calibration curve.^36^

### ELISA chip video recording and still image capture

For video recording (Panasonic Lumix DMC-GH3K), the ELISA chips were loaded with the ELISA buffer solution in the reagents and washing buffer inlets and with the 4×-diluted pooled saliva solution in the sample inlet. Both solutions were colored with food dye for visualization purposes unless stated otherwise.

Videos were edited in Adobe Premiere Pro (Version: 22.1.2) to adjust brightness, contrast, sharpness, and speed. Still images were captured using the Sony α7R III camera and edited in Adobe Photoshop for brightness, contrast, and sharpness.

### Characterization of ELISA chip aliquoting and maximum over-loading capacity

To characterize aliquoting, screenshots of the chips following the completion of aliquoting were analyzed in Fiji. The regions corresponding to the extra or lost solution shown in Fig. S3 were mapped, and the equivalent volume was calculated based on the 3D design file. The visual aliquoted volume and the nominal aliquoting accuracy were then calculated as follows:

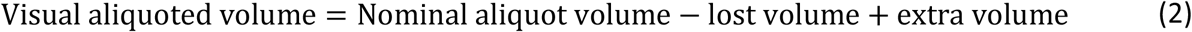

Where the nominal aliquot volume is equal to the capacity of the reservoir in the 3D design.

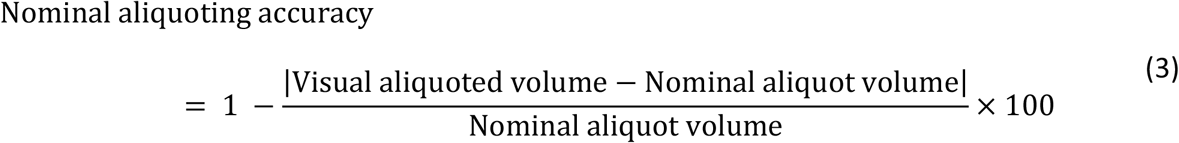

To characterize the maximum overloading capacity, the chips were loaded with exact volumes of sample, reagents and washing buffer using laboratory micropipettes with an increment of 5 µL for each solution. The recorded videos were analyzed visually to investigate any unwanted mixing or spilling of reagents to the adjacent reservoirs.

### Characterization of ELISA chip timing

The videos of the ELISA chips were recorded, and the timing of each step in the MCR read from the video data was tabulated and the duration of different steps was calculated.

### Capillary pressure and resistance calculation

Capillary pressure was calculated using the Young-Laplace equation as follows:^19^

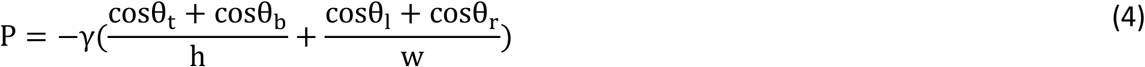

Where P denotes the capillary pressure, γ represents the liquid surface tension, h and w are the microchannel height and widths, and θ_t_, θ_b_, θ_l_, and θ_r_ respectively are the contact angle of the liquid with the top, bottom, left, and right wall of the microchannel.^19^ For the ELISA buffer and 4×-diluted saliva, the liquid surface tension γ was measured to be 37.8 and 38.0 mN·m^-1^ respectively using a dynamic tensiometer (DCAT11; Filderstadt; Germany). A custom-built apparatus was utilized for the contact angle measurement. Each θ_b_, θ_l_, and θ_r_ was replaced by the contact angle of ELISA buffer (10 ± 1.5°) or 4×-diluted saliva (12.5 ± 2°) on the plasma-treated, flat 3D samples. θ_t_ was replaced by the contact angle of the ELISA buffer (86.2 ± 3.4°) or 4×-diluted saliva (88.7 ± 1.9°) on the microfluidic diagnostic tape.

To calculate the resistance of a fluidic path, a lumped-element model was created where each section of the circuit was assigned a resistance calculated using the following equation:^19^

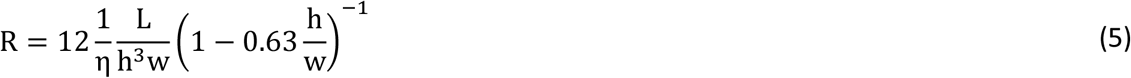

Where R denotes the resistance, η represents the liquid viscosity, and h is the height of the microchannel. Then, for circuit elements in series or parallel, the equivalent resistance (R_eq_) was calculated as follows:

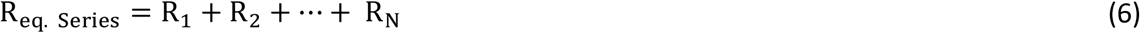

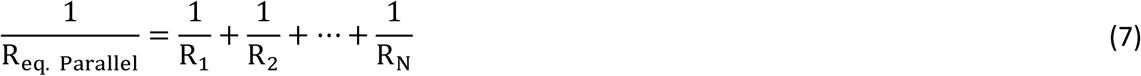

Where N denotes the number of capillaric elements included in the corresponding fluidic path. The liquid viscosity, η, was measured to be 1.11 mPa·s (temperature: 16.9 °C) for the ELISA buffer and 1.07 mPa·s (temperature: 17.7 °C) for the 4×-diluted saliva using a viscometer (SV-10; A&D Company Ltd; Tokyo; Japan).

## Authors contributions

A.P. and O.Y. designed the ELISA chip. A.P characterized the ELISA chip. A.P., J.R. W.J., and O.Y. optimized the SARS-CoV-2 N protein assay. A.P. and D. J. wrote the initial draft of the manuscript. A.P. and O. Y prepared the figures. A.P., O.Y., A.N., and D.J. reviewed and edited the manuscript. A.P. and DJ analyzed the data. D.J. conceptualized, administered, and supervised the work.

## Conflicts of interest

Authors declare that they have no conflicts of interest.

## Acknowledgments

We thank Geunyong Kim and Molly Shen for their assistance in assay optimization. We thank Mohamed Yafia for his constructive feedback regarding chip design. We thank Galyna Shul from NanoQAM, UQAM for her assistance in operating the viscometer and tensiometer. This work was supported by the NSERC Alliance Grant and the McGill MI4 Grant, and NSERC Discovery Grant. D.J. acknowledges support from a Canada Research Chair in Bioengineering.

## Supplementary Figures

**Fig. S1.**
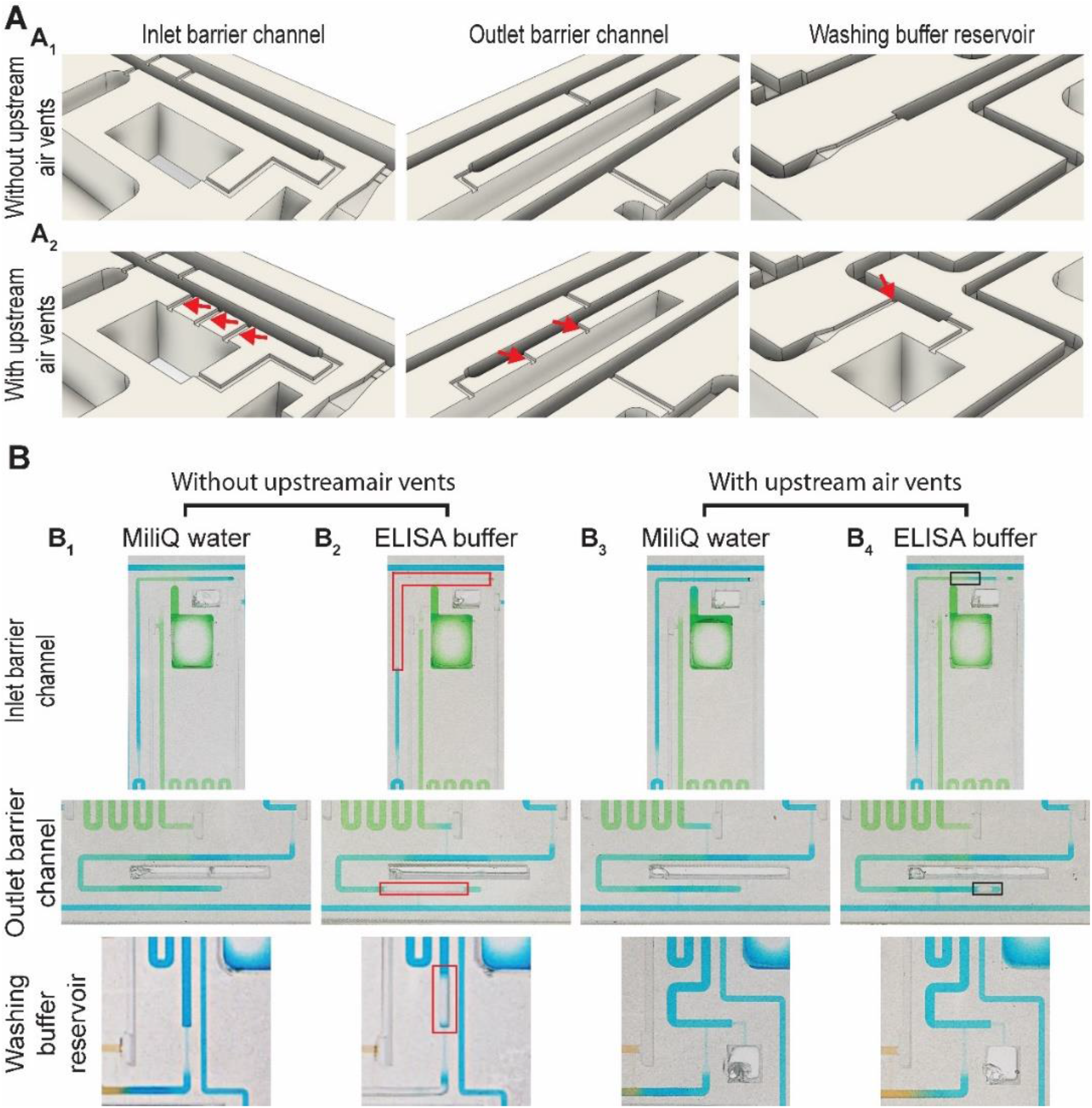
The effect of upstream air vents on the filling behavior of the ELISA chip. (A) The schematic illustration of the inlet barrier channel, outlet barrier channel, and the last washing buffer reservoir with and without upstream air vents, indicated by red arrows. (B) Filling with ELISA buffer containing 0.05% Tween 20 or MiliQ water in the absence or presence of upstream air vents. In the case of ELISA buffer with a low surface tension, the air vent at the channel end filled prematurely leading to a trapped air bubble (see the regions shown by the red rectangle in B1). Following the addition of upstream air vents, the conduits fill up to the desired point, even as a small bubble forms further downstream (see the black rectangles in B4). The solutions were colored with food dye for visualization purposes.

**Fig. S2.**
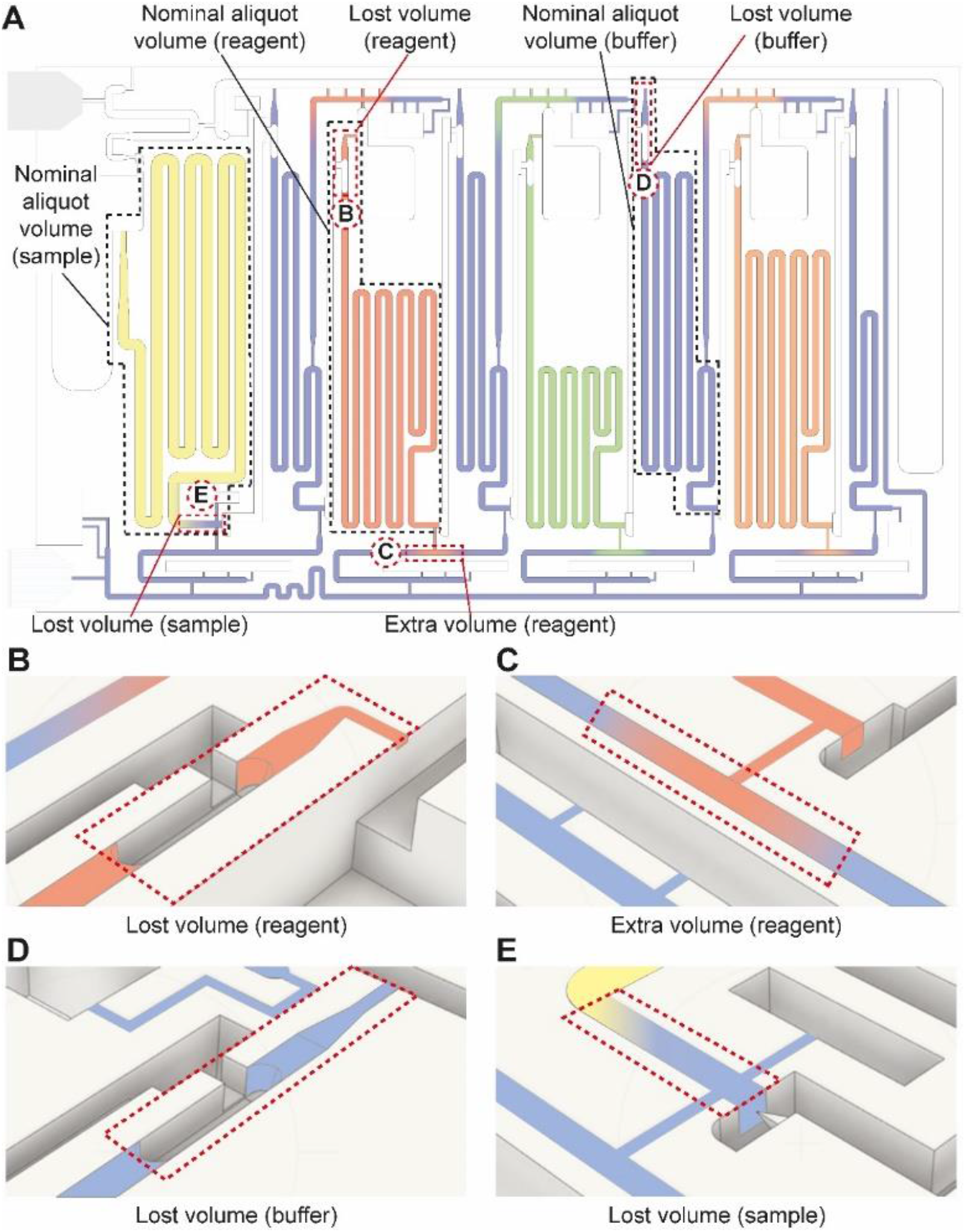
Illustration of lost and excess volumes in the ELISA chip following based on experimental observations. See video S1 for illustration of some of these examples. (A) The regions associated with the nominal aliquot volumes, lost, and extra volumes. (B-C) Close-up views of the lost and extra volumes in A. Note that lost volume could occur due to bubble formation in the reagent (See panel B) or washing buffer measuring reservoirs (See panel D), or due to overflow of the washing buffer to the sample measuring reservoir (See panel E), while extra volume could occur owing to the overflow of the reagents to the outlet barrier channel (See panel C).

**Fig. S3.**
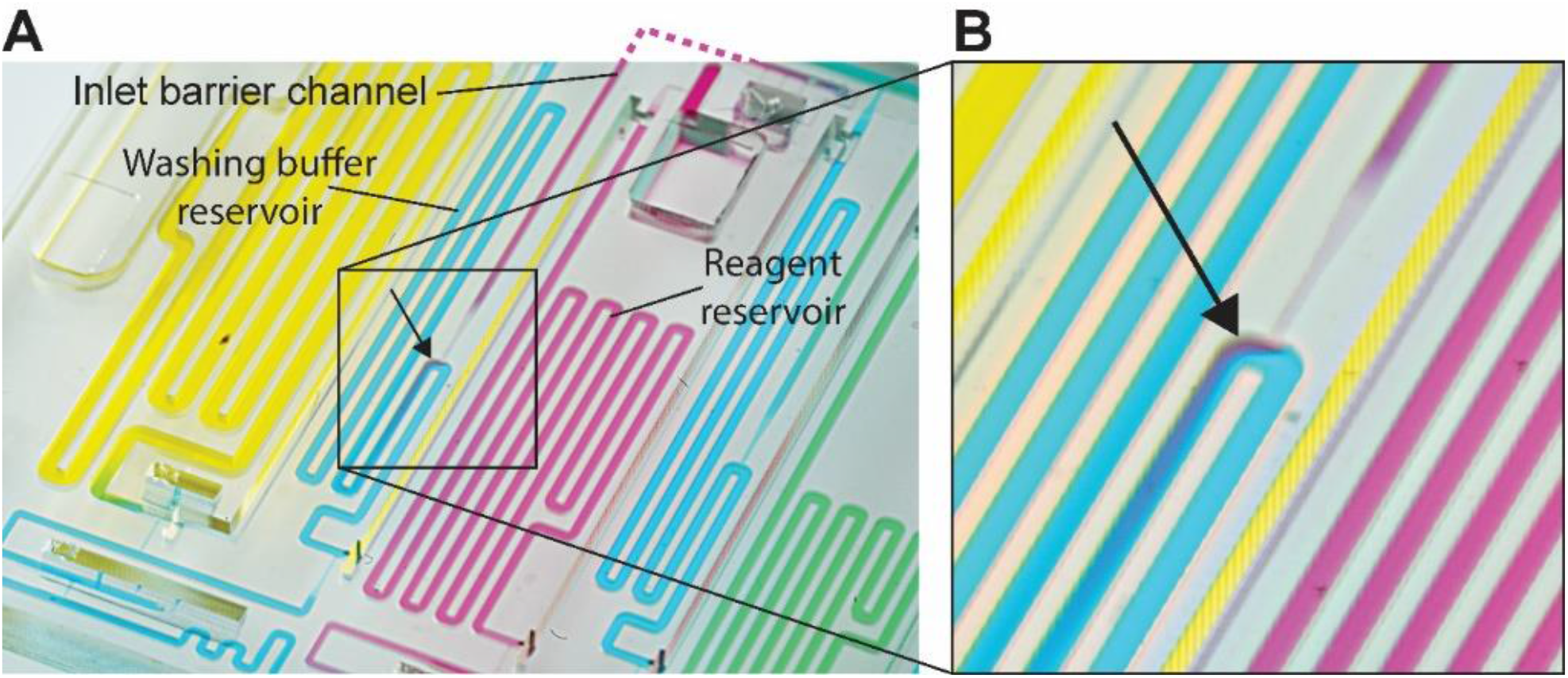
Reagent overlow in the ELISA chip. Photograph of the ELISA chip after loading the measuring reservoir with a nominal capacity of 70 µL with 140 µL of red reagent. The close-up photograph reveals the flow of the red-colored solution into the cyan buffer serpentine. The missing connecting channel that was outside of the frame of the picture was schematized using a dashed line.

**Fig. S4.**
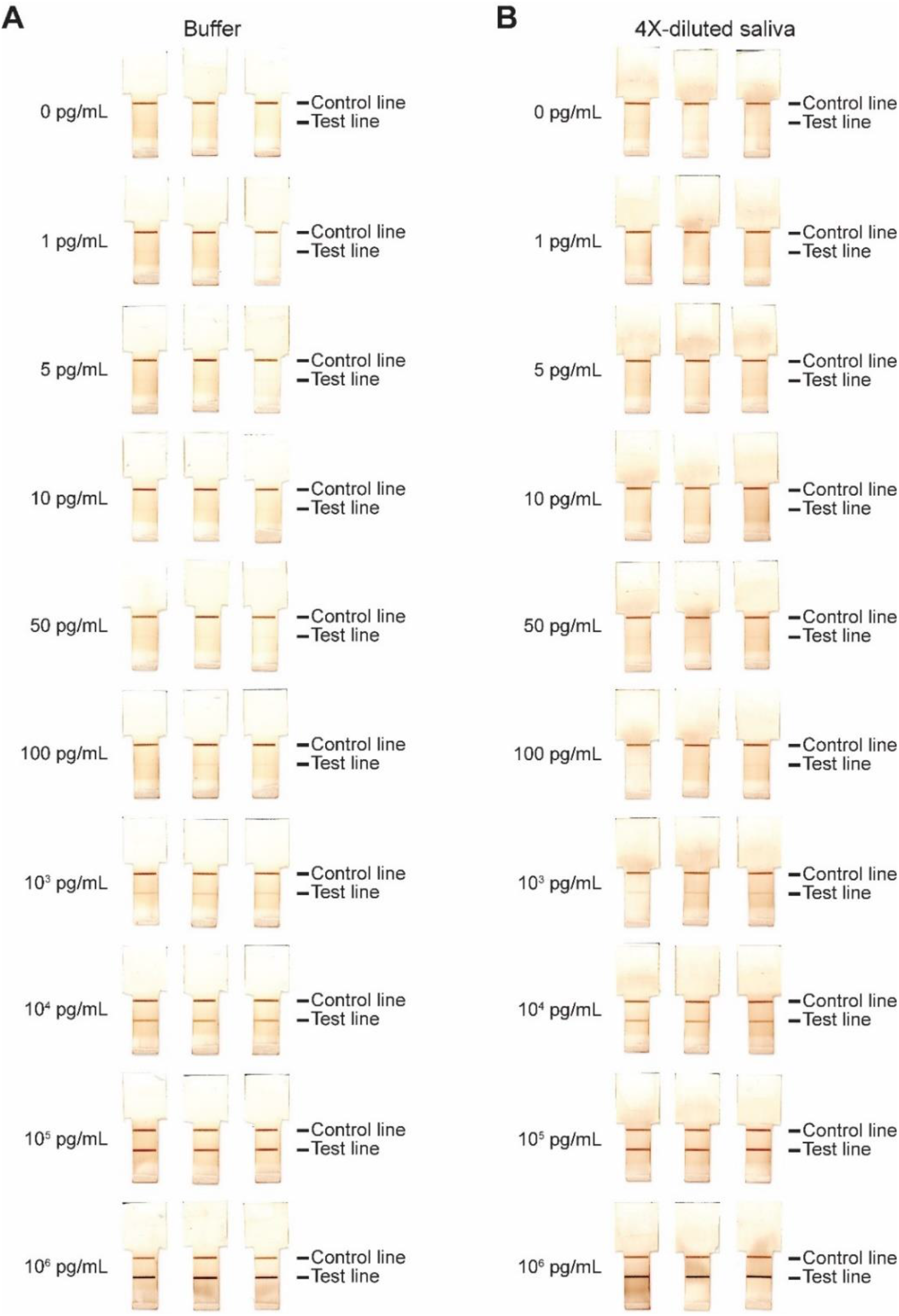
Scanned images of the nitrocellulose membranes for the SARS-CoV-2 N protein ELISA. Results of ELISA chip assays in (A) buffer and (B) 4×-diluted saliva for ten concentrations of spiked in N protein and the negative control, with three replicates each.

**Fig. S5.**
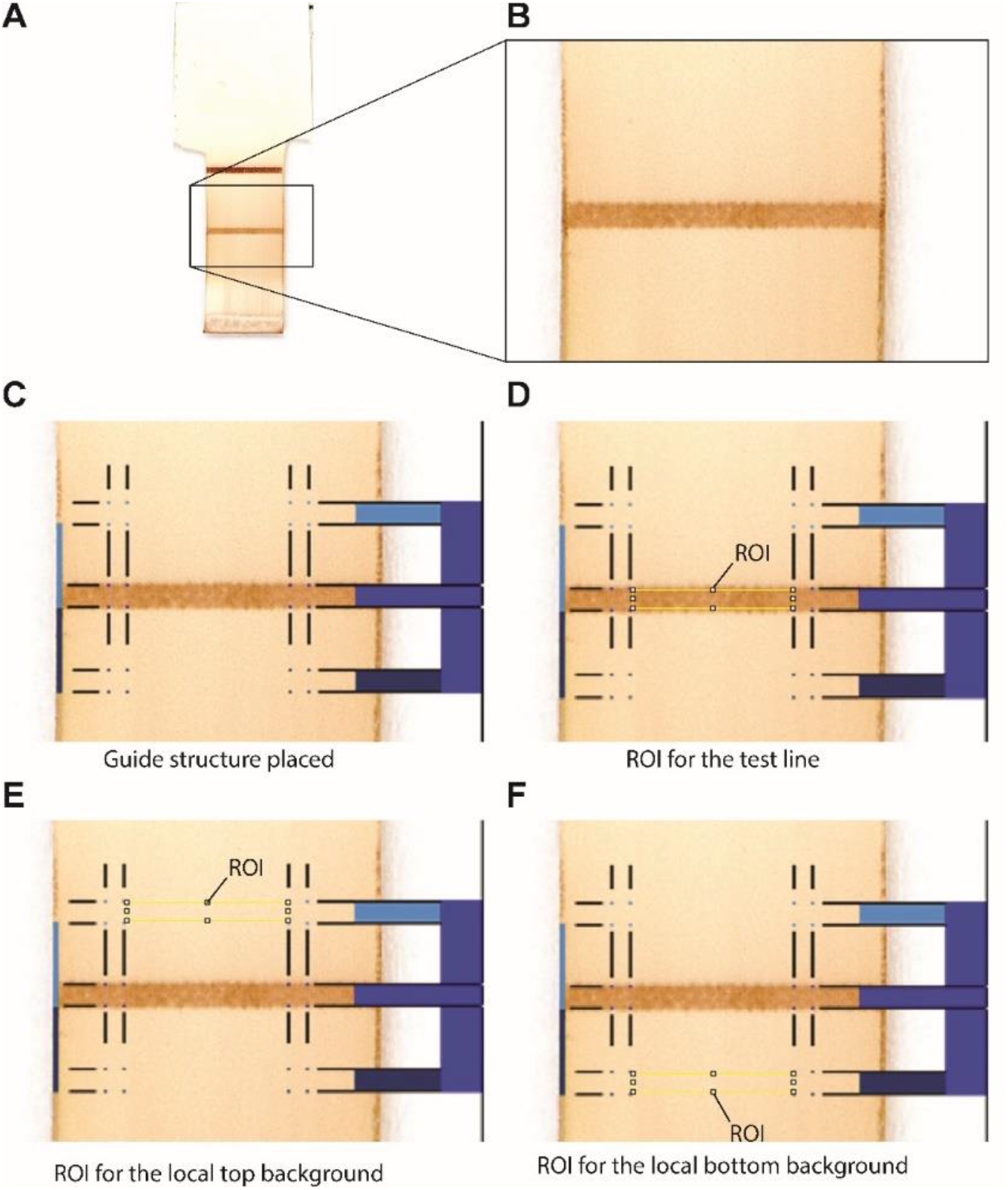
Image analysis of nitrocellulose membranes. (A) Scanned image of the nitrocellulose membrane. (B) Close-up view rectangle shown in panel A with the test line. (C) Nitrocellulose membrane superposed with the guide structure to position regions of interest (ROIs) to measure the signal intensity and background. (D – F) The ROIs for the test line as well as the top and bottom backgrounds used for gray value measurement.

## Supplementary tables

**Table S1.** Resistance of the entire path and the split section for each of paths i, ii, iii, iv.

| Fluidic path | Resistance of the entire path<sup>1</sup><br>(Pa·s·mm<sup>-3</sup>) | Resistance of split section<sup>2</sup><br>Value (Pa·s·mm<sup>-3</sup>) | Ratio<br>Path x/Path i |
| --- | --- | --- | --- |
| Path i | 135 | 18 | 7.8 |
| Path ii | 250 | 140 |  |
| Path i | 135 | 65 | 8.5 |
| Path iii | 620 | 550 |  |
| Path i | 135 | 95 | 6.9 |
| Path iv | 700 | 660 |  |

<sup>1</sup>This value was calculated for a given fluidic path (path i, ii, iii, or iv) starting from the reagent inlet and ending at the drainage capillary pump (See Fig. 3B).
<sup>2</sup>This value was calculated based on the point where a given parasitic path (path ii, iii, or iv) splits from drainage path i to the point where it merges again to path i (see Fig. 3B).

**Table S2.** Major steps of the SARS-CoV-2 N protein assay optimization.

| **Optimization** | **Range tested** | **Final condition** |
| --- | --- | --- |
| **Capturing antibody** | 0.07 - 0.28 µg / membrane | 0.28 µg /membrane |
| **Detection antibody** | 5 – 20 µg/mL | 7.5 µg/mL |
| **Enzyme conjugate** | 5 – 20 µg/mL | 7.5 µg/mL |
| **Tween 20** | 0.0125 – 0.1% | 0.05% |
| **Saliva dilution** | 1 – 10× | 4× |

## Notes

### Competing Interest Statement

The authors have declared no competing interest.

## References

1. S. D. Gan and K. R. Patel. Enzyme immunoassay and enzyme-linked immunosorbent assay. J. Invest. Dermatol. 2013, 133, e12.

2. Hosseini, P. Vázquez-Villegas, M. Rito-Palomares and S. O. Martinez-Chapa. Advantages, Disadvantages and modifications of conventional ELISA. SpringerBriefs Appl. Sci. Technol. Springer, Singapore, 2018, 67–115.

3. F. Costantini, C. Sberna, G. Petrucci, C. Manetti, G. de Cesare, A. Nascetti and D. Caputo. Lab-on-chip system combining a microfluidic-ELISA with an array of amorphous silicon photosensors for the detection of celiac disease epitopes. Sens. Bio-Sensing Res., 2015, 6, 51–58.

4. M. J. Uddin, N. H. Bhuiyan and J. S. Shim. Fully integrated rapid microfluidic device translated from conventional 96-well ELISA kit. Sci. Reports, 2021, 11, 1–9.

5. L. Zhu, Y. Feng, X. Ye, J. Feng, Y. Wu and Z. Zhou. An ELISA chip based on an EWOD microfluidic platform. J. Adhes. Sci. Technol., 2012, 26, 2113–2124.

6. Ng, A. H. C. et al. A digital microfluidic system for serological immunoassays in remote settings. Sci. Transl. Med. 10, (2018).

7. J. T. Heggestad et al. Multiplexed, quantitative serological profiling of COVID-19 from blood by a point-of-care test. Sci. Adv., 2021, 7, 4901–4926.

8. S. Laschi, R. Miranda-Castro, E. González-Fernández, I. Palchetti, F. Reymond, J. S. Rossier, G. Marrazza and S. Laschi. A new gravity-driven microfluidic-based electrochemical assay coupled to magnetic beads for nucleic acid detection. Electrophoresis, 2010, 31, 3727– 3736.

9. N. M. Reis, S. H. Needs, S. M. Jegouic, K. K. Gill, S. Sirivisoot, S. Howard, J. Kempe, S. Bola, K. Al-Hakeem, I. M. Jones and T. Prommool. Gravity-driven microfluidic siphons: fluidic characterization and application to quantitative Immunoassays. ACS Sensors, 2021, 6, 4338–4348.

10. K. M. Koczula and A. Gallotta. Lateral flow assays. Essays Biochem., 2016, 60, 111–120.

11. G. A. Posthuma-Trumpie, J. Korf and A. Van Amerongen. Lateral flow (immuno)assay: Its strengths, weaknesses, opportunities and threats. A literature survey. Anal. Bioanal. Chem., 2009, 393, 569–582.

12. Y. Liu, L. Zhan, Z. Qin, J. Sackrison and J. C. Bischof. Ultrasensitive and highly specific lateral flow assays for point-of-care diagnosis. ACS Nano, 2021, 15, 3593–3611.

13. C. Parolo, A. de la Escosura-Muñiz and A. Merkoçi. Enhanced lateral flow immunoassay using gold nanoparticles loaded with enzymes. Biosens. Bioelectron., 2013, 40, 412–416.

14. Y. Panraksa, I. Jang, C. S. Carrell, A. G. Amin, O. Chailapakul, D. Chatterjee, C. S. Henry. Simple manipulation of enzyme-linked immunosorbent assay (ELISA) using an automated microfluidic interface. Analytical Methods, 2022, 14, 1774–1781.

15. M. S. Verma, M. N. Tsaloglou, T. Sisley, D. Christodouleas, A. Chen, J. Milette and G. M. Whitesides. Sliding-strip microfluidic device enables ELISA on paper. Biosens. Bioelectron., 2018, 99, 77–84.

16. M. M. Gong and D. Sinton. Turning the Page: Advancing Paper-Based Microfluidics for Broad Diagnostic Application. Chem. Rev., 2017, 117, 8447–8480.

17. B. J. Toley, J. A. Wang, M. Gupta, J. R. Buser, L. K. Lafleur, B. R. Lutz, E. Fu and P. Yager. A versatile valving toolkit for automating fluidic operations in paper microfluidic devices. Lab Chip, 2015, 15, 1432–1444.

18. Safavieh, R. & Juncker, D. Capillarics: pre-programmed, self-powered microfluidic circuits built from capillary elements. Lab Chip 13, 4180–4189 (2013).

19. R. Safavieh and D. Juncker. Capillarics: pre-programmed, self-powered microfluidic circuits built from capillary elements. Lab Chip, 2013, 13, 4180–4189.

20. O. Gökçe, S. Castonguay, Y. Temiz, T. Gervais and E. Delamarche. Self-coalescing flows in microfluidics for pulse-shaped delivery of reagents. Nature, 2019, 574, 228–232.

21. T. U. Vinitha, S. Ghosh, A. Milleman, T. Nguyen, C. H. Ahn. A new polymer lab-on-a-chip (LOC) based on a microfluidic capillary flow assay (MCFA) for detecting unbound cortisol in saliva. Lab Chip, 2020, 20, 1961–1974.

22. E. E. Ahi, H. Torul, A. Zengin, F. Sucularlı, E. Yıldırım, Y. Selbes, Z. Suludere and U. Tamer. A capillary driven microfluidic chip for SERS based hCG detection. Biosens. Bioelectron., 2022, 195, 113660.

23. N. Ramalingam, H. B. Liu, C. C. Dai, Y. Jiang, H. Wang, Q. Wang, K. M. Hui, H. Q. Gong. Real-time PCR array chip with capillary-driven sample loading and reactor sealing for point-of-care applications. Biomed. Microdevices, 2009, 11, 1007–1020.

24. M. Yafia, O. Ymbern, A. O. Olanrewaju, A. Parandakh, A. Sohrabi Kashani, J. Renault, Z. Jin, G. Kim, A. Ng and D. Juncker. Microfluidic Chain Reaction of Structurally Programmed Capillary Flow Events. Nature, 2022, 605, 464–469.

25. M. Steinitz. Quantitation of the Blocking Effect of Tween 20 and Bovine Serum Albumin in ELISA Microwells. Anal. Biochem., 2000, 282, 232–238.

26. P. B. Luppa, C. Müller, A. Schlichtiger and H. Schlebusch. Point-of-care testing (POCT): Current techniques and future perspectives. TrAC Trends Anal. Chem., 2011, 30, 887–898.

27. V. Gubala, L. F. Harris, A. J. Ricco, M. X. Tan and D. E. Williams. Point of care diagnostics: Status and future. Anal. Chem., 2012, 84, 487–515.

28. S. Sachdeva, R. W. Davis and A. K. Saha. Microfluidic point-of-care testing: Commercial landscape and future directions. Front. Bioeng. Biotechnol., 2021, 8, 1537, p.602659

29. C. D. Chin, T. Laksanasopin, Y. K. Cheung, D. Steinmiller, V. Linder, H. Parsa, J. Wang, H. fMoore, R. Rouse, G. Umviligihozo and E. Karita. Microfluidics-based diagnostics of infectious diseases in the developing world. Nat. Med., 2011, 17, 1015–1019.

30. Z. He, J. Huffman, K. Curtin, K. L. Garner, E. C. Bowdridge, X. Li, T. R. Nurkiewicz and P. Li. Composable microfluidic plates (cPlate): A simple and scalable fluid manipulation system for multiplexed enzyme-linked immunosorbent assay (ELISA). Anal. Chem., 2021, 93, 1489–1497.

31. M. Jing, R. Bond, L. J. Robertson, J. Moore and A. Kowalczyk. User experience of home-based AbC-19 SARS-CoV-2 antibody rapid lateral flow immunoassay test. Sci. Reports, 2022, 12, 1–18.

32. C. C. Miller. The Stokes-Einstein law for diffusion in solution. Proc. R. Soc. London. Ser. A, Contain. Pap. a Math. Phys. Characte., 1924, 106, 724–749.

33. E. S. Savelaz et al. Quantitative SARS-CoV-2 viral-load curves in paired saliva and nasal swabs inform appropriate respiratory sampling site and analytical test sensitivity required for earliest viral detection. J. Clin. Microbiol., 2022, 60, e01785–21.

34. V. M. Corman et al. Comparison of seven commercial SARS-CoV-2 rapid point-of-care antigen tests: a single-centre laboratory evaluation study. The Lancet Microbe, 2021 2, 311–319.

35. A. De Lean, P. J. Munson and D. Rodbard. Simultaneous analysis of families of sigmoidal curves: application to bioassay, radioligand assay, and physiological dose-response curves. Am. J. Physiol. Endocrinol. Metab., 1987, 4, E:97.

36. C. K. Dixit, S. K. Vashist, B. D. MacCraith and R. O’Kennedy. Multisubstrate-compatible ELISA procedures for rapid and high-sensitivity immunoassays. Nat. Protoc., 2011, 6, 439– 445.

